# Movement patterns and activity levels are shaped by the neonatal environment in Antarctic fur seal pups

**DOI:** 10.1101/2021.03.09.434640

**Authors:** Rebecca Nagel, Sina Mews, Timo Adam, Claire Stainfield, Cameron Fox-Clarke, Camille Toscani, Roland Langrock, Jaume Forcada, Joseph I. Hoffman

## Abstract

Tracking studies of juveniles are rare compared to those of adults, and consequently little is known about the influence of intrinsic and extrinsic factors on activity during this critical life stage. We used hourly GPS data, collected from 66 Antarctic fur seal pups from birth until moulting, to investigate the explanatory power of multiple individual-based and environmental variables on activity levels. Pups were sampled from two nearby breeding colonies of contrasting density during two subsequent years, and a two-state hidden Markov model was used to identify modalities in their movement behaviour, specifically ‘active’ and ‘inactive’ states. We found that movement was typified by central place exploration, with active movement away from and subsequent return to a location of inactivity. The probability of such directed exploration was unaffected by several factors known to influence marine mammal movement including sex, body condition, and temperature. Compared to pups born at the high-density colony, pups at low-density were more active, increased their activity with age, and transitioned earlier into the tussock grass, which offers protection from predators and extreme weather. Our study illustrates the importance of extrinsic factors, such as colony of birth, to early-life activity patterns and highlights the adaptive potential of movement.

## Introduction

Movement is a defining characteristic of life that underpins critical components of behaviour^1^ and fitness^2^. Understanding how animals alter their movement in response to external and internal stimuli is, therefore, fundamental to the management and conservation of wild populations^1,3^. Rapid advancements in bio-logging technology and the increased accessibility of statistical tools for drawing meaningful inference from fine-scaled observations have generated unprecedented insights into the movement of a variety of species in their natural habitats^4,5^. Marine vertebrates in particular have benefited from such improved methodologies, exemplified by a recent analysis of more than 2,600 tracked individuals documenting extraordinary convergence in movement patterns across 50 species^6^.

Despite these recent advancements in marine tracking research, several authors have drawn attention to age and sex biases in the literature, with datasets of adult females being particularly over-represented^3,7^. This focus on adults is problematic because the movement and distribution of neonatal and juvenile individuals is of key importance for understanding population dynamics. For example, long-term monitoring of three albatross species has shown that high rates of juvenile mortality due to interactions with fisheries are likely driving observed declines in population size^8^. Similarly, juvenile and adult loggerhead sea turtles occupy distinct oceanic environments, making young turtles more susceptible to bycatch on pelagic longlines^9^.

In addition to informing conservation efforts, studies of the movement of young individuals can also provide valuable insights into the ontogeny of social and survival skills. A tracking study of European shags, for example, has shown that higher rates of juvenile mortality correlate with poor foraging proficiency^10^, highlighting the importance of learning and memory for successful recruitment^11^. Studies of several different pinniped species, including Antarctic fur seals^12^, New Zealand sea lions^13^, Steller sea lions^14^, northern fur seals^15^, and grey seals^16^ also suggest that habitat choice as adults may be primarily driven by intrinsic factors early in life rather than size-related differences as adults.

These and other studies represent an important step forward in the field of movement ecology because they help to close the gap between adult movement and the ‘lost years’ of juveniles^3^. However, with few exceptions, studies of juveniles have focused on the period after nutritional independence. While this can be justified for many birds and other species where neonates are more or less stationary until weaned, in many other species movement behaviour and social interactions among conspecifics earlier in life play a key role in development. For example, play behaviour in pre-weaned Stellar sea lions and Galápagos fur seals facilitates the development of muscle mass and hones fighting skills that are important for future foraging and reproductive success^17,18^.

Antarctic fur seals (*Arctocephalus gazella*) are another pinniped species where neonatal movement may play an important role in early development. Newborn pups are regularly left unattended in dense rookeries while their mothers, who are central place foragers, intersperse short periods of nursing on land with longer foraging trips at sea^19^. Without their mothers to defend them, pups are at an increased risk of predation from predatory birds^20^ and traumatic injury due to crushing by territorial males^21,22^. Furthermore, the natal coat of pups lacks the water-repellent properties of adult fur^23^, which prevents them from spending prolonged time at sea. Consequently, pups can only begin to develop the efficient swimming and diving behaviour necessary for nutritional independence after they moult at around 60 days of age^24,25^. Anecdotal observations of pups venturing into shallow pools and streams shortly after birth^24^ and the emergence of sex-specific differences in habitat use later in life^26^ provide some insights into pup activity, but we still know surprisingly little about the intrinsic and extrinsic factors that influence pup movement prior to moulting.

Here, we analysed hourly GPS data from 66 Antarctic fur seal pups tracked from birth until moulting (Fig. 1) using a hidden Markov model. Focal individuals were selected at random from two breeding colonies on Bird Island, South Georgia, the Special Study Beach (SSB) and Freshwater Beach (FWB), which are separated by around 200 meters (Fig. 2a). Because of their close proximity, these colonies experience comparable climatic conditions and breeding females from both locations likely forage in the same area^27^, which is reflected by the fact that females do not differ significantly in quality traits such as body size or condition^28^. Both colonies also offer comparable cobblestone substrate habitats backed by tussock grass, which covers most of the island’s interior and offers protection from predators and extreme weather. Despite these similarities, the density of animals is higher at SSB than at FWB^28^, which creates a different dynamic at the two colonies. First, tightly packed males at SSB have little freedom to move, which results in a more static system^29^, and second, because avian predators can more readily penetrate low density breeding aggregations, there is a greater likelihood of pups being predated on FWB^28^. The two colonies also differ in their overall area and topology. SSB is somewhat smaller and is relatively ‘closed’ as a steep gully separates the beach from the tussock grass immediately inland. By contrast, FWB is larger and has flatter terrain making it more ‘open’.

**Figure 1:**
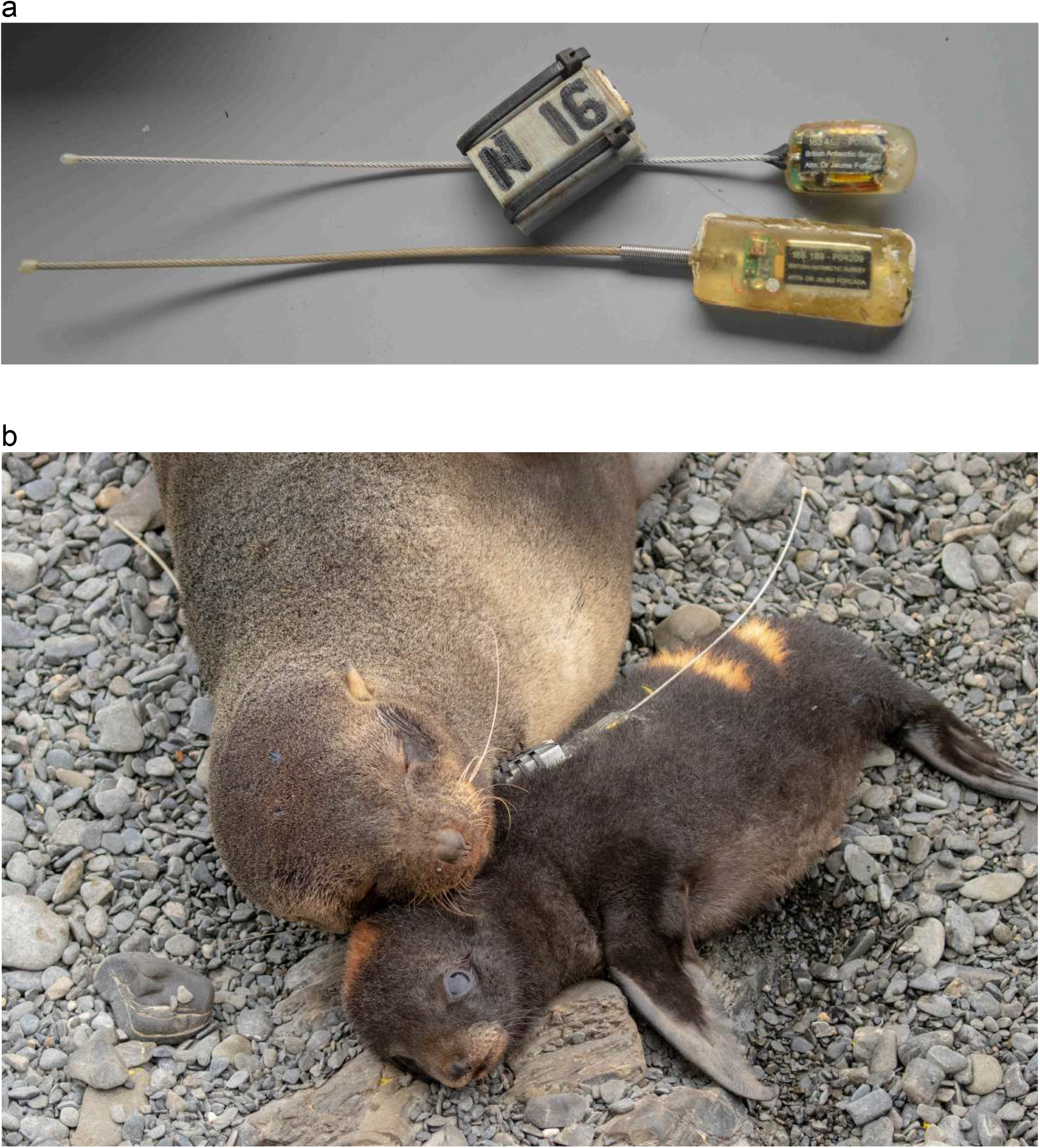
VHF monitoring and GPS tracking. (a) VHF transmitter and GPS logger for pups (top) and VHF transmitter for mothers (bottom). (b) Antarctic fur seal mother-pup pair. VHF transmitters are visible on both individuals, while the pup has additionally been fitted with a GPS logger and given a temporary bleach mark for identification. Photo credit: Claire Stainfield.

**Figure 2:**
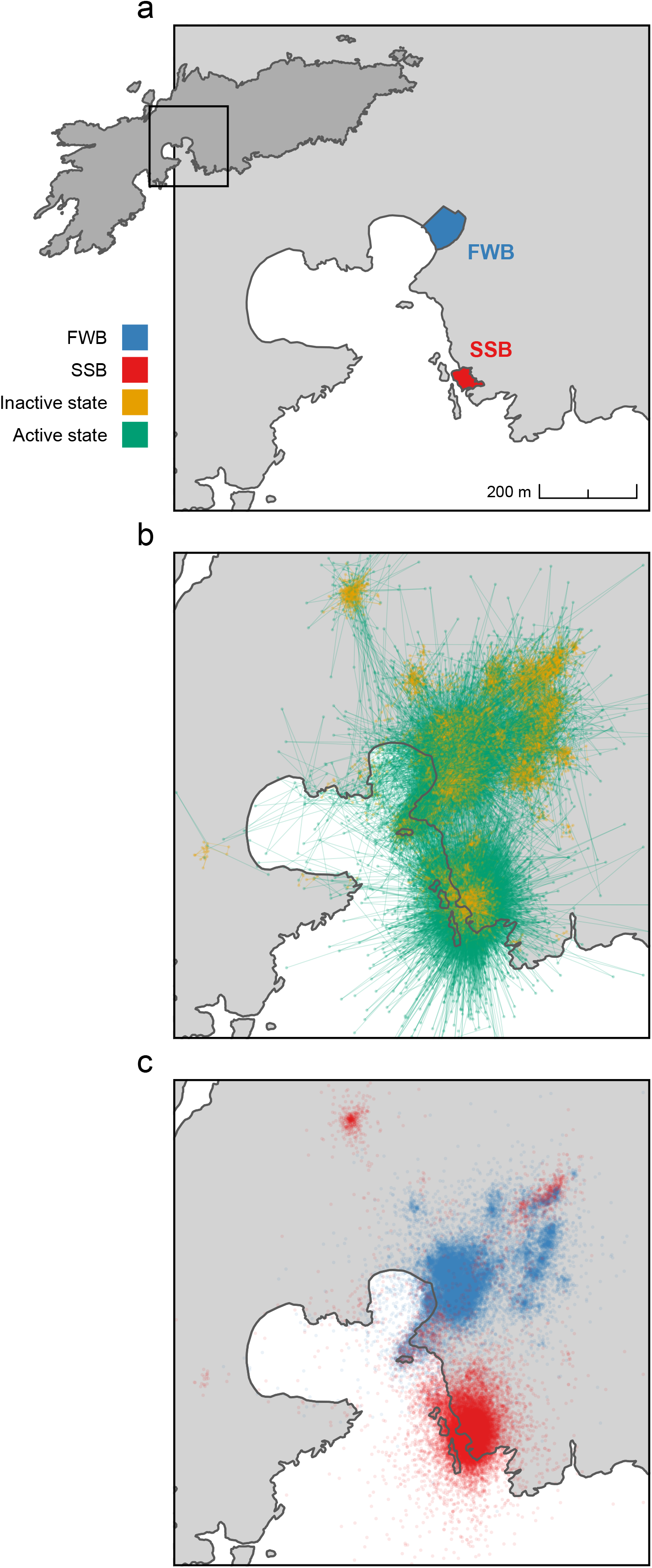
Movement patterns of Antarctic fur seal pups on Bird Island, South Georgia. (a) Location of the two fur seal breeding colonies, Freshwater Beach (FWB) and the Special Study Beach (SSB), from which pups were tagged with GPS loggers. (b) Decoded states from the HMM showing the inferred activity patterns of 66 pups throughout ontogeny. (c) GPS locations recorded throughout ontogeny for 32 pups born on FWB and 34 pups born on SSB. To aid visibility, each panel is a magnified view of Bird Island (longitude range = −38.060 - 38.045, latitude range = −54.014 −54.005) and consequently, a small proportion of decoded states and GPS points recorded outside this frame are not visible.

We used these two colonies as a ‘natural experiment’ to investigate the influence of the early neonatal environment on the movement of fur seal pups, as well as to evaluate the relative importance of intrinsic (e.g. sex, age, and body condition) versus extrinsic (e.g. colony) factors. Furthermore, as the number of breeding individuals, foraging trip durations, and pup birth weights vary appreciably from year to year depending on local food abundance^30^, we replicated our study across two consecutive breeding seasons of contrasting food availability^28^. We hypothesised that (i) intrinsic and extrinsic factors shown in previous studies to influence movement in other pinnipeds, including sex^31–33^, body condition^34^, and ambient temperature^35,36^, affect the movement of Antarctic fur seal neonates, (ii) activity levels and movement patterns are influenced by density, with FWB pups being more active to avoid predation, and (iii) levels of pup activity increase with age, as there is likely strong selection for the early development of motor skills and lean muscle mass.

## Results

Hourly GPS data were successfully collected from a total of 66 Antarctic fur seal pups from two nearby breeding colonies of contrasting density (Fig. 2a). Deployment durations ranged from 20 to 80 days (median = 51 days) for the 53 surviving pups and from 4 to 41 days (median = 15 days) for the 13 pups that died. On average, pups travelled 43.9 meters per hour. In general, pup movement showed a star-like pattern characterised by directed exploration within a relatively small area around a central location of low activity (Fig. 2b). However, the spatial distribution of these ‘home patches’ varied by colony. Pups born at FWB remained significantly closer to their natal colony (Wilcoxon rank sum test, *p* = 0.002), traveling on average 87.6 meters into the tussock grass immediately inland. By contrast, pups born at SSB moved an average of 205.0 meters from their natal colony in a wider variety of directions (Fig. 2c). For a map of the entire island showing all recorded positions, see Supplementary Fig. S1. For representative examples of individual GPS tracks, including one pup from each breeding colony and year, see Supplementary Fig. S2.

### Hidden Markov model: Activity patterns of fur seal pups

The two states of the HMM were clearly discriminated from each other. State 1 captured smaller step lengths (mean step length = 22.5 meters) corresponding to less active movement and hereafter referred to as the ‘inactive state’. State 2 was characterised by more active behaviour covering longer distances (mean step length = 75.8 meters) and hereafter referred to as the ‘active state’. Minimal differences between the marginal distribution under the fitted model and the empirical distribution as well as the analysis of pseudo-residuals suggest that our model provides a good fit to the data (see Supplementary Fig. S3 for more details). Based on the Viterbi-decoded state sequences, pups spent on average 65.2% (range = 45.7 – 97.3%) of their time in the inactive state and 34.8% (2.7 – 54.3%) of their time in the active state. The pups’ decoded movement patterns are summarised in Fig. 2b.

Differences in AIC values (delta AIC) between the full model and HMMs sequentially excluding each variable and its interaction term with colony are shown in Table 1. We considered variables to be relevant for modelling pup activity levels if their delta AIC values were considerably larger than zero (≥ 40). Based on this criterion, air temperature, wind speed, sex, and body condition had little to no influence on activity levels. By contrast, the covariates year, time of day, and age were determined to have the strongest effects on pup activity levels via both main effects and interactions with colony (Table 1). Pups tended to be slightly more active in the first year of the study, particularly at FWB where the probability of occupying the active state fell from 43.8% in 2019 to 34.0% in 2020. Activity levels also showed a clear diurnal pattern, peaking in the middle of the day in both colonies (Fig. 3a and b). Finally, marked differences were observed in colony-specific developmental trajectories, with animals from FWB exhibiting an almost linear increase in activity with age (Fig. 3c), whereas activity levels at SSB remained more or less constant throughout ontogeny (Fig. 3d). FWB pups also tended to venture inland into the tussock grass earlier than pups from SSB and spent proportionally more time in the tussock grass than in their colony of birth (Fig. 3e and f).

**Table 1:**
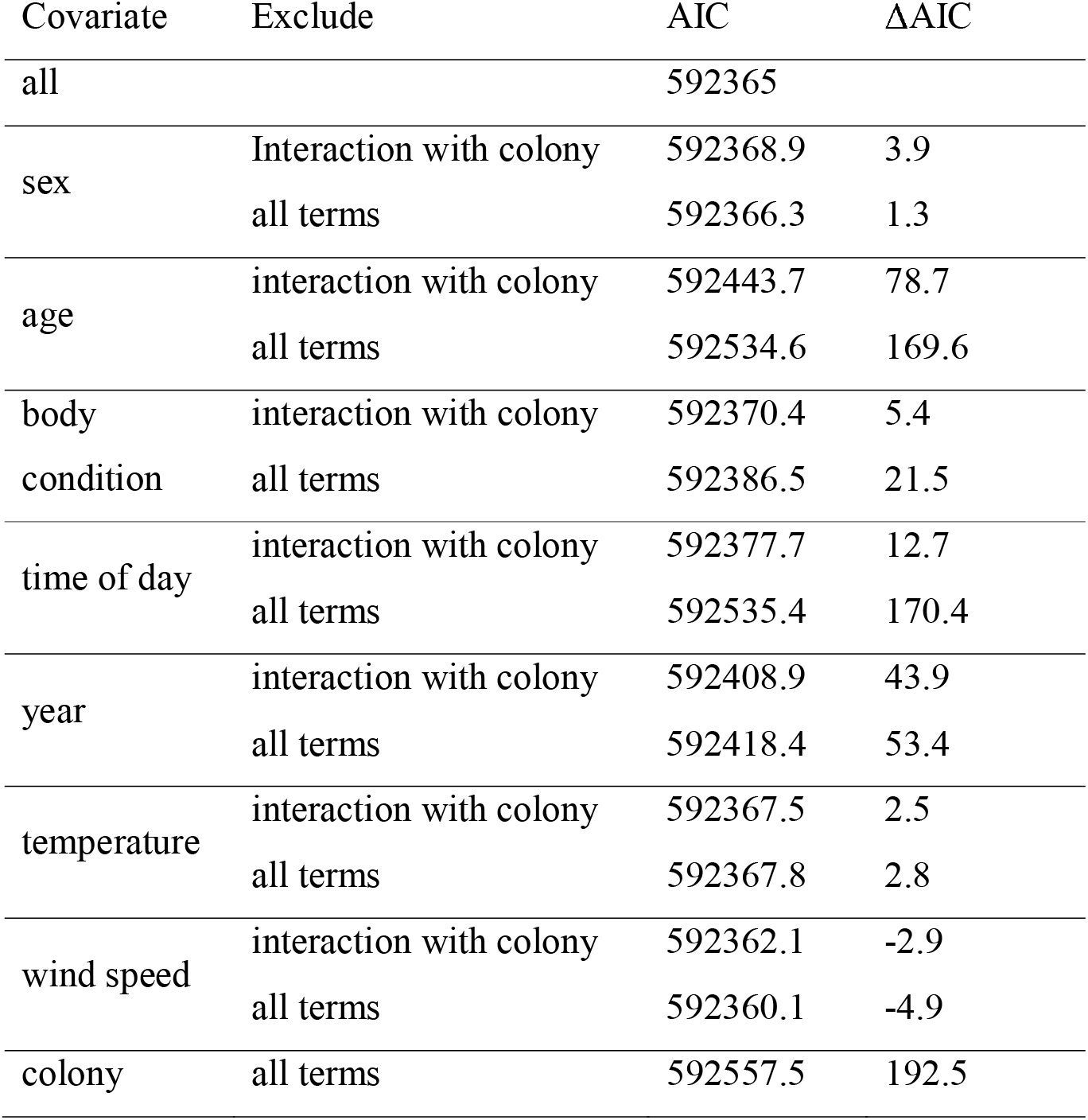
Differences in AIC values (ΔAIC) between the full model and HMMs sequentially excluding each variable and its interaction term with colony.

**Figure 3:**
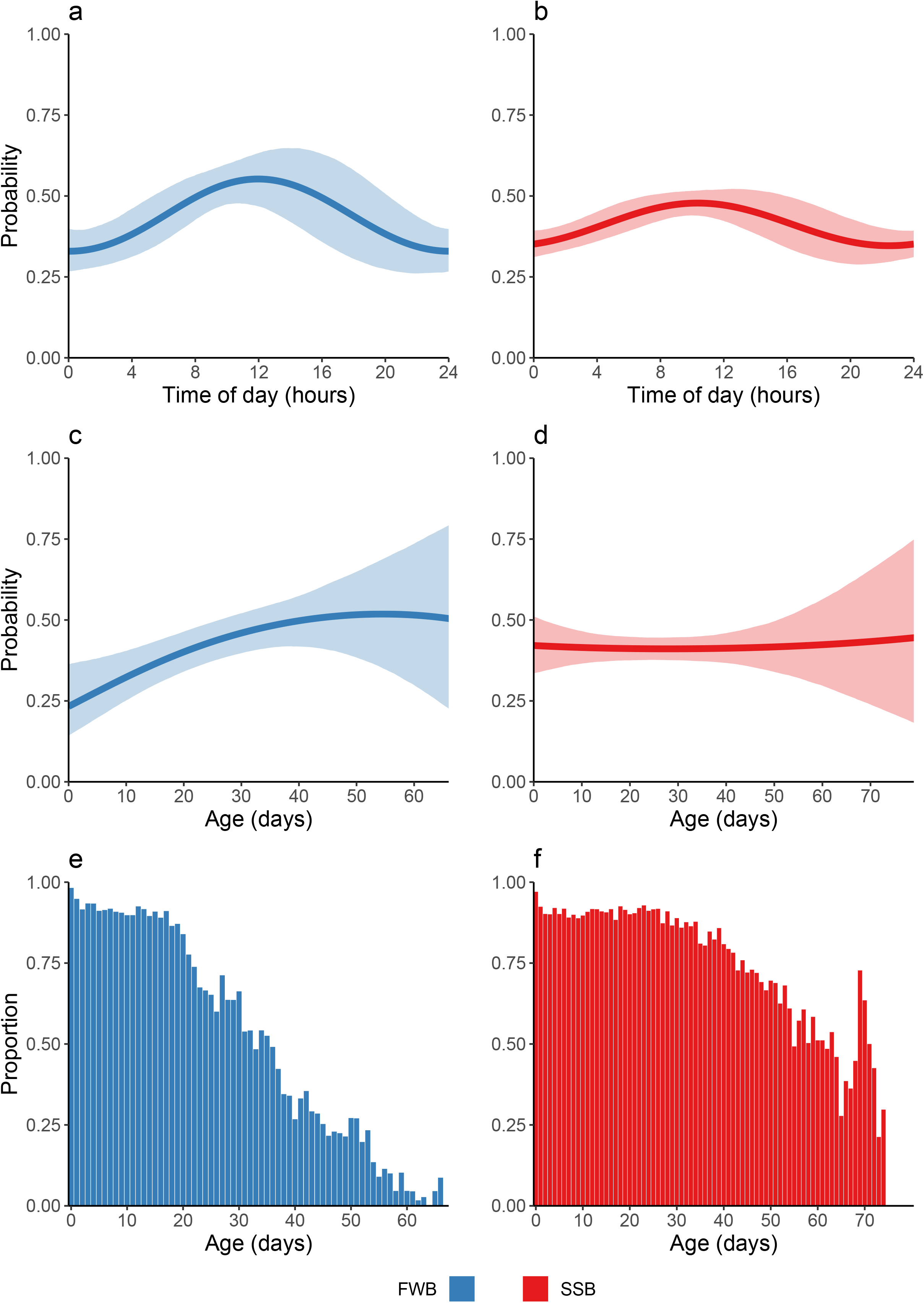
Main results of the HMM. (a–d) Mean probability (± 95% confidence interval) of being in an ‘active’ behavioural state as a function of the covariates included in the HMM. Probabilities were calculated for each covariate and state by fixing the other covariates to their respective means or reference categories. Confidence intervals for the probabilities were obtained based on Monte Carlo simulation from the estimators’ approximate distribution as implied by maximum likelihood theory. (a, b) The probably of being in an active state as a function of time. (c, d) The probability of being in an active state as a function of age. (e, f) The proportion of time pups spent at their colony of birth or elsewhere on the island.

### Post-hoc analyses

To test additional hypotheses that could not be addressed in the HMM, we performed *post-hoc* logistic regression analyses to evaluate whether the presence of the mother ashore and the fate of the pup (died or survived until the end of the study) explained a significant proportion of the variation in pup activity levels (’states’). Pups were significantly more active (8%) when their mothers were absent from Bird Island on foraging trips compared to when their mothers were present (Fig. 4a; GLMM, estimate = 0.09, *s.e.* = 0.04, *p*-value = 0.03). Pups that survived were also significantly more active (52%) than pups that died (GLMM, estimate = 0.758, *s.e.* = 0.19, *p*-value < 0.001) (Fig. 4b).

**Figure 4:**
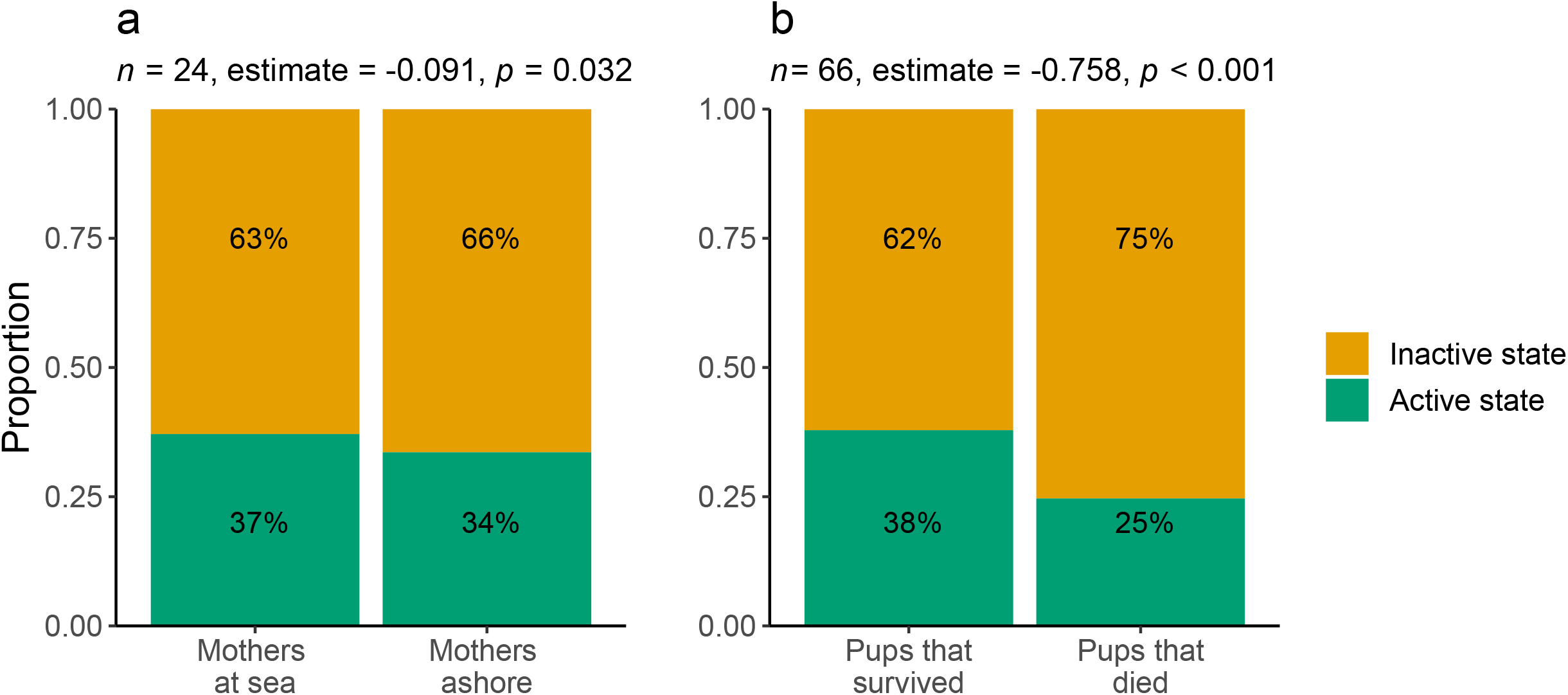
Results of *post-hoc* analyses. Proportion of time pups spent in the active and inactive states (a) when their mothers were at sea foraging or ashore nursing and (b) depending on mortality status. Sample sizes (*n*), GLMM slope estimates, and *p*-values are provided.

## Discussion

We collected and analysed hourly GPS data from Antarctic fur seal pups tracked from birth until moulting in order to investigate the drivers of movement patterns and activity levels during this critical life stage. We found that pup movement was characterised by a star-like pattern, with individuals repeatedly returning to a central location of low activity after bouts of directed exploration. The HMM showed that the probability of such active movement was highest during the day and was mainly influenced by breeding season, colony of birth, and age. Our findings provide new insights into the movement patterns of pinnipeds prior to nutritional independence and highlight the importance of the life-history stage and the neonatal environment on behaviour.

### Movement in fur seal pups

A recent study of movement in Antarctic fur seal pups found evidence for sex-specific differences in habitat use prior to nutritional independence and the onset of overt sexual size dimorphism. Specifically, male pups were found to exhibit more risk prone behaviour with increasing age, traveling further at sea and spending more time in exposed beach habitats^26^. While this study highlighted the importance of an intrinsic factor, sex, on neonatal movement, the contributions of other factors, both extrinsic and intrinsic, remained largely unexplored. Our HMM incorporated multiple environmental and individual-based variables including the time of day, air temperature, wind speed, age, body condition, and sex. Rather than focusing on animals from a single colony, our study used tracking data from two breeding colonies with contrasting population density and terrain, while replication across two consecutive breeding seasons also allowed us to account for inter-annual variation in food availability.

Our hourly GPS data revealed a distinct pattern of terrestrial pup movement that has not been reported in previous studies. Specifically, we documented a star-like pattern characterised by directed movement away from and subsequent return to a location of low activity. This appears to be a genuine behavioural pattern rather than an artefact of our temporal scale of sampling, as GPS locations collected every five minutes from a subset of individuals revealed a very similar picture (Supplementary Fig. S4). Star-like movement patterns are typified by central place foragers, which make round trips between a central location and a foraging patch^37^. However, as fur seal pups are entirely reliant on their mothers for food until they moult, repeated movements away from a central location are unlikely to be related to resource acquisition. Instead, our results appear to be indicative of bouts of exploration within a defined radius around a suckling location, which would imply that Antarctic fur seal pups are ‘central place explorers’.

We would expect central place exploration in fur seal pups to be adaptive given that it entails energetic costs^37^ and might even make it more difficult for pups to be located by their mothers when they return from foraging trips^38^. We can envisage a number of possible explanations for this behaviour. First, high levels of activity may facilitate the development of the muscle mass necessary for future foraging success. Second, movement both within and beyond the confines of the beach where a pup is born may increase the scope for social interactions, which might be important for future mating success^18^. Third, a transition from beach habitats to the tussock grass offers increased protection against harsh weather conditions and may reduce the risk of a pup either being crushed by a territorial male or predated.

While all tracked individuals displayed this pattern of central place exploration, the distance travelled during bouts of exploration differed depending on the colony of birth. On average, pups born at FWB travelled shorter distances and stayed closer to their colony of birth, whereas SSB pups moved farther and in a larger variety of directions. Given that SSB pups transition into the tussock grass later in life (see discussion below), one explanation for this difference may be that mother-pup pairs need to travel further inland to find an appropriate, unoccupied suckling location.

### Results of the HMM

We used an HMM framework to infer the discrete behavioural states underlying pup movement and the probability of switching between these states given certain environmental and individual covariates. This integrated approach to decompose behavioural patterns has been applied across a wide range of species and habitats and has emerged as a powerful tool for animal movement modelling^5^. Our model uncovered a strong influence of year, with the probability of occupying the active state being higher in the first breeding season (2019), particularly for FWB pups. One possible explanation for this pattern could be that inter-annual differences in movement are adaptive. The first breeding season in our study had among the lowest breeding female numbers, pup birth weights, and foraging trip durations on record^28^, reflecting poor environmental conditions and low prey abundance^39^. In 2019, local densities at both colonies were correspondingly below average^28^. Given a greater risk of predation at low density^28^, pups might increase their activity to avoid harassment, pecking injuries, and predation by generalist seabirds such as southern and northern giant petrels^40^. Alternatively, higher levels of activity could be a consequence of increased stress hormone levels. As mothers spent more time at sea foraging in 2019^28^, pups were subjected to longer periods of food deprivation. Such extended bouts of starvation in pinniped pups have been associated with an increase in glucocorticoids^41^, which are known to increase activity levels in some species^42^. Understanding the interrelationships among prolonged fasting, glucocorticoids levels, and activity in fur seal pups may help to shed further light on individual responses to environmental challenges.

Another clear extrinsic determinant was the time of day, with activity levels peaking just before solar noon. This observation is in line with a previous study of Galápagos fur seal pups^18^, but otherwise very little is known in general about diurnal patterns of activity among fur seals while ashore. Daily meter-resolution location data from densely packed territory holding Antarctic fur seal males reveal negligible movements from day to day^29^, implying that these animals remain more or less stationary. By contrast, telemetry studies of adult females have reported associations between nocturnal foraging behaviour and diel variation in the time of arrival and departure from the breeding colony^43^. Consequently, it is unclear to what extent our results can be extrapolated across the life history, especially given the likelihood of an ontogenetic shift in times of peak activity as pups transition from being reliant on milk to nutritional independence.

The HMM also revealed an increase in pup activity with age, but only for pups born at FWB. One explanation for this colony-specific difference could be that FWB pups are able to express their full behavioural repertoire throughout the course of ontogeny, becoming more active as they grow, whereas pups born on SSB are constrained by the high density of conspecifics. In particular, pups that traverse tightly packed harems on SSB run a higher risk of being crushed by a territorial male or bitten by a breeding female^21,22^. Alternatively, we recently found that FWB pups are more likely to be predated by generalist seabirds^28^. This might translate into an increase of predator-avoidance activity as pups mature, which would help to explain why activity no longer increases after around 40 days of age, when the majority of individuals have transitioned to the more sheltered tussock grass.

We found that pups from FWB tended to move into the tussock grass earlier than pups from SSB and spent proportionally more time in the tussock grass than in their colony of birth. While this earlier shift toward residing mainly in the tussock grass could simply be a consequence of increased activity, it might also be adaptive for FWB pups to move inland as quickly as possible given the higher risk of predation at low density^28^. In other words, pups may move into the more sheltered tussock grass earlier in life and remain there for longer periods of time in order to avoid predatory seabirds. However, differences in the topography of the two colonies might also influence the timing of this transition, as a steep gully separates the beach at SSB from the tussock grass, while FWB offers a more gradual transition between the two habitats. To disentangle the effects of density and topography would require a larger study embracing a greater diversity of breeding colonies.

Contrary to our initial expectations, the HMM found little to no effect of air temperature or wind speed on pup activity. This is surprising given that the pups in our study lacked the water-repellent fur of adults and were thus poorly protected from the elements^23^. Reduced activity^35^ and huddling behaviour^44^ have been documented in pinniped neonates as effective thermoregulatory behaviours to withstand the respectively hottest and coldest daily temperatures, so we originally anticipated a reduction in pup activity under marginal weather conditions. However, large variation in temperature during the course of this study was not observed on Bird Island (2019 mean temperature = 3.7°C ± *s.d*. 1.3; 2020 mean temperature = 3.9°C ± *s.d*. 1.4). Consequently, behavioural adjustments in activity for effective thermoregulation may not have been necessary in the context of our study. Alternatively, other climatic variables that we could not account for, such as precipitation or humidity, may have a disproportionate influence on activity levels. Future studies involving the direct observation of focal individuals under specific climatic conditions would help to address this question.

The HMM also showed that pup activity levels were largely unaffected by body condition. This was unexpected because Antarctic fur seal pups rely on their mother’s milk for nutrition before moulting and must tolerate bouts of starvation lasting up to 11 days while their mothers forage at sea^19^. As a result, pup growth is known to decline with prolonged maternal absence^45^, implying that fewer resources should be available for movement. Taken at face value, the lack of a relationship between body condition and movement may suggest that the short-term benefits of high activity, such as muscle mass development and increased social interaction, may outweigh the costs associated with diverting resources from growth. This would be in line with a previous study that found that Galápagos fur seal pups maintained high activity levels throughout bouts of starvation^18^. However, it is also possible that an effect of body condition could not be detected in our study because our model assumed that condition was constant between successive measurements, whereas in practice it will vary to some extent from day to day.

Despite a number of studies having shown that sex-specific differences in activity, habitat use, foraging, and diving behaviour are established early in life in several pinniped species^13–17,46^, including Antarctic fur seals^26^, our HMM did not uncover any obvious sex differences in activity. This is probably a consequence of the timeframe of our study. Jones *et al.*^26^, for example, only detected sex-specific differences in habitat use in Antarctic fur seal pups greater than 41 days old. Given that we focused on the time window from birth until moulting at around 60 days of age, the results of these two studies are consistent and lend support to the notion that sex-specific movement patterns take several weeks to become established.

### *Results of the* post-hoc *analysis*

The *post-hoc* analysis showed that pups were significantly less active when their mothers were ashore, suggesting that pup activity is correlated to some extent with maternal foraging behaviour. This association is most likely a reflection of the utmost importance of milk consumption for pup survival and development. A mother may spend as little as 24 hours and on average only two days ashore during each nursing bout^43^, so pups must maximise nutrient update during this time. Moreover, adult female fur seals frequently display aggressive behaviour towards foreign pups^21^, potentially limiting a pup’s opportunity for social interactions when the mother is present^18^. It is important to note, however, that although significant, the effect size of the correlation between pup activity and maternal presence was fairly small. This suggests that pups may not remain with their mothers for the entire maternal attendance period, but rather alternate between nursing and bouts of activity.

We also found that pups that died tended to be considerably less active. In our dataset, cause of death was assigned to all but one pup as either starvation (*n* = 5) or predation (*n* = 6), two factors that are difficult to untangle given that smaller, weaker (i.e. starving) pups may be more likely to be predated. It is possible that the reduced activity of pups that died could be related to poor body condition. While this would appear to contradict the lack of an overall association between activity and condition in the HMM, it is possible that activity may decline only after body condition falls below a critical threshold^18^, an effect we may not have captured due to the small number (*n* = 13) and early occurrence (median = 15 days) of mortalities in our dataset. Alternatively, causality could flow in the other direction, with less active pups being more likely to be predated. This would be more in line with our HMM results as well as with the hypothesis that increased activity is related to predator-avoidance behaviour.

### Strengths, limitations, and future directions

Recent advancements in bio-logging technology and analytical methods have made tracking studies of small, juvenile individuals ethically feasible, cost effective, and analytically approachable. In this study, we have taken advantage of these advancements to implement one of the first in-depth analyses of the movement and activity patterns of Antarctic fur seal pups prior to moulting. We were able to uniquely incorporate individual, ecological, and environmental variation into our analyses by collecting time-series biometric data for all tagged individuals, which were sampled at two breeding colonies across two subsequent breeding seasons. Finally, we were able to tease apart how these intrinsic and extrinsic factors may influence movement by inferring behavioural states using a HMM.

An important limitation of our study, however, is our inability to establish causal relationships between hypothesised explanatory factors and activity. While this is an ever-present challenge when working with wild populations, future studies may consider pairing GPS tracking data with detailed behavioural observations of the focal individuals. This would provide a behavioural context for movement patterns and possibly allow for a more nuanced interpretation of the underlying behavioural states. A higher GPS sampling rate might also facilitate the resolution of finer-scale states. While this was not possible in the current study due to the limited battery capabilities of our tags, as technologies continue to improve future studies may not need to sacrifice sampling resolution for study length. Finally, it would be interesting in the future to consider GPS tracking both mothers and pups to better understand the relationship between maternal attendance behaviour and pup activity. This approach might also shed light on how mothers find their offspring after returning ashore from foraging trips.

### Conclusions

We document the movement behaviour and activity levels of Antarctic fur seal pups between birth and moulting in relation to a multitude of extrinsic and intrinsic explanatory variables. Our findings suggest that during this early life stage, pup activity is mainly shaped by extrinsic factors including year, time of day, and colony of birth. In contrast to a previous study of activity later in life^26^, we found little effect of intrinsic factors such as body condition and sex. Our study highlights the importance of external influences during this critical phase of life when pups are at their greatest risk of predation and have not yet developed their water-repellent adult coat.

## Methods

### Field methods

This study was conducted at two breeding colonies on Bird Island, South Georgia (54°00’24.8□ S, 38°03’04.1□ W; Fig. 2a) during the austral summers (December to March) of 2018 – 19 (hereafter referred to as 2019) and 2019 – 20 (hereafter referred to as 2020).

In both breeding seasons, 25 Antarctic fur seal pups from each colony were captured two to three days after birth (December) and every ten days thereafter until they began to moult (March). Individuals were randomly selected with respect to sex and birth date during the pupping period. Capture, restraint, and handling of pups followed protocols established over 36 consecutive years of the long-term monitoring and survey program of the British Antarctic Survey (BAS). Briefly, pups were captured with a slip noose or by hand and were restrained by hand. After handling, pups were returned to their mothers or released as close to their capture sites as possible. At every capture, weight and length measurements were taken, from which a scaled mass index was calculated according to Peig and Green^47^. This condition metric serves as a reliable indicator of overall fitness as it has been correlated with mortality^48^ and reproductive success^49^ in a variety of species.

To facilitate the tracking and recapture of focal individuals, VHF transmitters (Sirtrack core marine glue-on V2G 152A; dimensions: 40 × 20 × 10 mm body with a 200 mm antenna, weight: 16 g) were attached to the dorsal side of the neck between the shoulder blades with epoxy glue. Pups were similarly fitted with GPS loggers in waterproof ABS plastic enclosures (Perthold Engineering CatLog Gen2 GPS Loggers; dimensions: 50 × 40 × 17 mm, weight: 36 g), which recorded latitude and longitude positions within five-meter accuracy every hour from birth until moulting of the pelage (Fig. 1). Together, these two tags accounted for less than 1% of the mean focal pup mass at first capture (5.6 kg). Temporary bleach marks applied to the fur were used to identify pups. Unrecovered tags (*n* = 12) and bleach marks are shed naturally when the seals moult (March – April), precluding any long-term consequences for the pups.

As part of the BAS contribution to the Ecosystem Monitoring Programme of the Convention for the Conservation of Antarctic Marine Living Resources (CCAMLR), the attendance behaviour of breeding females has been monitored since 1982, with radio telemetry protocols established in 1992 to track around 25 adult females per year on FWB. We contributed towards this ongoing effort by attaching VHF transmitters (Sirtrack core marine glue-on V2G 154C; dimensions: 65 × 28 × 13 mm body with a 250 mm antenna, weight: 42 g) to the mothers of our focal FWB pups. Adult females were captured with a noosing pole and held on a restraint board. Daily attendance was monitored using a fixed-position radio antenna (Televilt RX900) combined with visual checks of the island with a hand-held VHF receiver (AOR Ltd., AR8200). The daily absence or presence of females ashore was noted from first capture until the final measurement, when the pups either moulted or died. Mothers were fitted with cattle ear tags in the trailing edge of each fore flipper for identification.

### Data analysis framework

Within our dataset, hourly GPS data were successfully collected from a total of 66 Antarctic fur seal pups (breakdown by year: *n* = 40 in 2019 and *n* = 26 in 2020; by colony: *n* = 32 at FWB and *n* = 34 at SSB). The majority of tracked individuals (*n* = 53, or 80%) survived until weaning. For those individuals that died, we truncated the GPS data back to the last time point that the animal was seen alive prior to analysis.

Our GPS data contained some missing locations, which in a handful of cases led to observation gaps of several hours. Most if not all missing values likely arose due to signal lapse, which occurs when the satellite connection to a GPS device is interrupted due to cloud cover, physical obstruction, etc.^50^. In an otherwise regular sequence, such missing data are manageable, but large gaps may introduce bias into the model^51^. Therefore, for observational gaps exceeding four hours, we split the individual’s track into separate bursts or intervals of continuous data. Bursts that were shorter than 48 hours usually contained a considerable amount of missing observations and were excluded from the analysis, corresponding to roughly 2% of the full dataset including missing locations. Overall, the tracks of 35 individuals were split into two to five bursts, while data from all of the other individuals (*n* = 31) consisted of a single burst.

Our final dataset contained 67,417 hourly observations with a median number of 1,124 observations per individual (min = 88, max = 1,916). The step lengths (in meters) between consecutive GPS locations were calculated and screened for implausible movements. From personal observations, we know that pups can move up to 400 meters in an hour. To confirm that 400 meters represents an appropriate step threshold, we analysed five-minute interval GPS data collected from three pups, each between 30 – 50 days of age, for 13 – 19 days (mean = 3,965 data points). In total, we found 2,029 instances where pups travelled a cumulative distance of more than 400 meters in an hour. To ensure these movements were biologically relevant and not due to erroneous recordings, we plotted a random subset and checked for unrealistic movement patterns, which were not present (see Supplementary Fig. S5). Based on these findings, we set all values larger than 400 meters/hour (corresponding to 297 observations) to be missing.

To investigate environmental factors influencing pup movement, hourly dry bulb air temperature and wind speed measurements for Bird Island were obtained from the BAS Meteorology and Ozone Monitoring Programme and missing values (85 observations) were linearly interpolated. The binary variables sex (with female as the reference category) and breeding season (with 2019 as the reference category) were held constant for each individual. The continuous variables age (measured in days since initial capture, two to three days after birth) and scaled mass index (calculated every ten days) were kept constant between days and measurements. All metric variables were standardised by subtracting the mean of the variable and dividing by its standard deviation.

### Hidden Markov models

To identify different activity states and investigate the explanatory power of environmental and individual-based variables on pup movement, we fitted a hidden Markov model (HMM) to the hourly GPS data. This time series model encompasses both the observed movement of an individual and an underlying (‘hidden’) state sequence, which is used to infer behavioural processes (e.g. active and inactive)^5^. HMMs have accordingly been used to analyse animal tracking data in relation to, among others, environmental conditions^52^, anthropogenic activity^53^, sex^54^, ontogeny^55^, and individualised niches^56^.

To distinguish between different behavioural processes in Antarctic fur seal pups, we fitted HMMs to the hourly observed step lengths. We assumed the observation process to be the same for all individuals and modelled the step lengths using a gamma distribution, conditional on the states. Prior to building a final model, we conducted exploratory analyses on the number of states and compared univariate HMMs to bivariate HMMs additionally including the turning angle between consecutive GPS locations. We restricted our final analysis to a parsimonious univariate HMM with two states for two reasons. First, the inclusion of the turning angle resulted in a negligible improvement of model fit. Second, the addition of a third state resulted in overlapping distributions of the step length, suggesting no additional information was gained while complicating the interpretation of the underlying states (see Supplementary Fig. S6)^57^. To check whether the HMMs based on hourly data adequately reflect the animals’ movement behaviour, we also fitted an HMM to the five-minute interval GPS data. The results showed that, with coarser temporal resolution (hourly data), there was no relevant loss of information on pup activity with respect to the aims of the study (see Supplementary Fig. S4 and S7).

We further investigated the effects of external and internal factors, including colony, breeding season, time of day, temperature, wind speed, sex, age, and body condition, on the activity level of fur seal pups by modelling state transition probabilities as a function of the covariates using a logit link function. We allowed the effects of the covariates to differ between the two colonies by including interaction terms with the binary covariate ‘colony’ (FWB/SSB). To account for the periodic nature of time of day, its effect was modelled using trigonometric functions. To assess the relative importance of the covariates, we calculated the differences in AIC values between the full model and models sequentially excluding each covariate and its interaction with colony. All HMMs were fitted by numerically maximising the likelihood using *moveHMM*^51^. To further investigate model fit, we calculated pseudo-residuals and checked them for normality and autocorrelation (Supplementary Fig. S3)^5^. Based on the full model, we inferred the most likely underlying state sequence using the Viterbi algorithm^5^. We also calculated the state occupancy probabilities as a function of each covariate^58^, while fixing the other covariates to their respective means or reference categories.

### Post-hoc analyses

The HMM decomposed the GPS data into two distinct states, which we inferred as ‘inactive’ and ‘active’ movement. To determine whether additional variables that we were not able to include in the HMM explain a significant proportion of variation in activity, we built two generalised linear mixed models (GLMMs) *post-hoc*. In our first model, we included maternal attendance (0 = present ashore, 1 = absent) as the explanatory variable. As data on maternal attendance were only available for pups born on FWB, this covariate could not be included in the full HMM, which defined an interaction term between all covariates and ‘colony’. In our second model, we used the survival status of each pup (0 = died, 1 = survived) as the explanatory variable. Here, we wanted to test for an association between survival status and movement without assuming the direction of causality. In both models, pup ID was included as a random effect to account for repeated measurements of individuals. The response variable was the inferred state of an individual (0 = inactive, 1 = active) at a given time point and was modelled with a binomial error distribution using *lme4*^59^. The residuals of the models were visually inspected for linearity and equality of error variances (using plots of residuals versus fits), normality (using Q – Q plots), and over or under-dispersion (by comparing the dispersion of simulated to observed residuals) using *DHARMa*^60^. All analyses and visualisations for this study were implemented in R version 4.0.2^61^.

## Animal ethics and permits

Animal handling was carried out by BAS under permits from the Government of South Georgia and the South Sandwich Islands (Wildlife and Protected Areas Ordinance (2011), RAP permit numbers 2018/024 and 2019/032). All procedures used were approved by the BAS Animal Welfare and Ethics Review Body (AWERB applications 2018/1050 and 2019/1058) and performed in accordance with provided guidelines and regulations.

## Data Availability Statement

Raw data and R code, available as a R Markdown file, have been uploaded to the Dryad Digital Repository (https://datadryad.org/stash/share/j6KfNHr2FNMlcw4TXOgcYbplq1E1gDa9xonh365a8Vc) and will be made public upon acceptance of the manuscript.

## Acknowledgements

The authors would like to thank Ana Bertoldi Carneiro, Freya Blockley, Jamie Coleman, Alexandra Dodds, Vicki Foster, Derren Fox, Iain Angus Gordon, Pauline Goulet, Rosie Hall, Cary Jackson, Adam Lowndes, Elizabeth Morgan, Rachael Orben, Jessica Ann Philips, David Reid, and Mark Whiffin for help in the field.

## Author contributions

RN, CS, CF-C, and CT collected the data. SM and TA built the HMM models. JIH, RL, and JF conceived of the study and contributed funding and materials. All of the authors commented on and approved the final manuscript.

