## Supplementary Material for "Movement patterns and activity levels are shaped by the neonatal environment in Antarctic fur seal pups"

**Supplementary Figures**

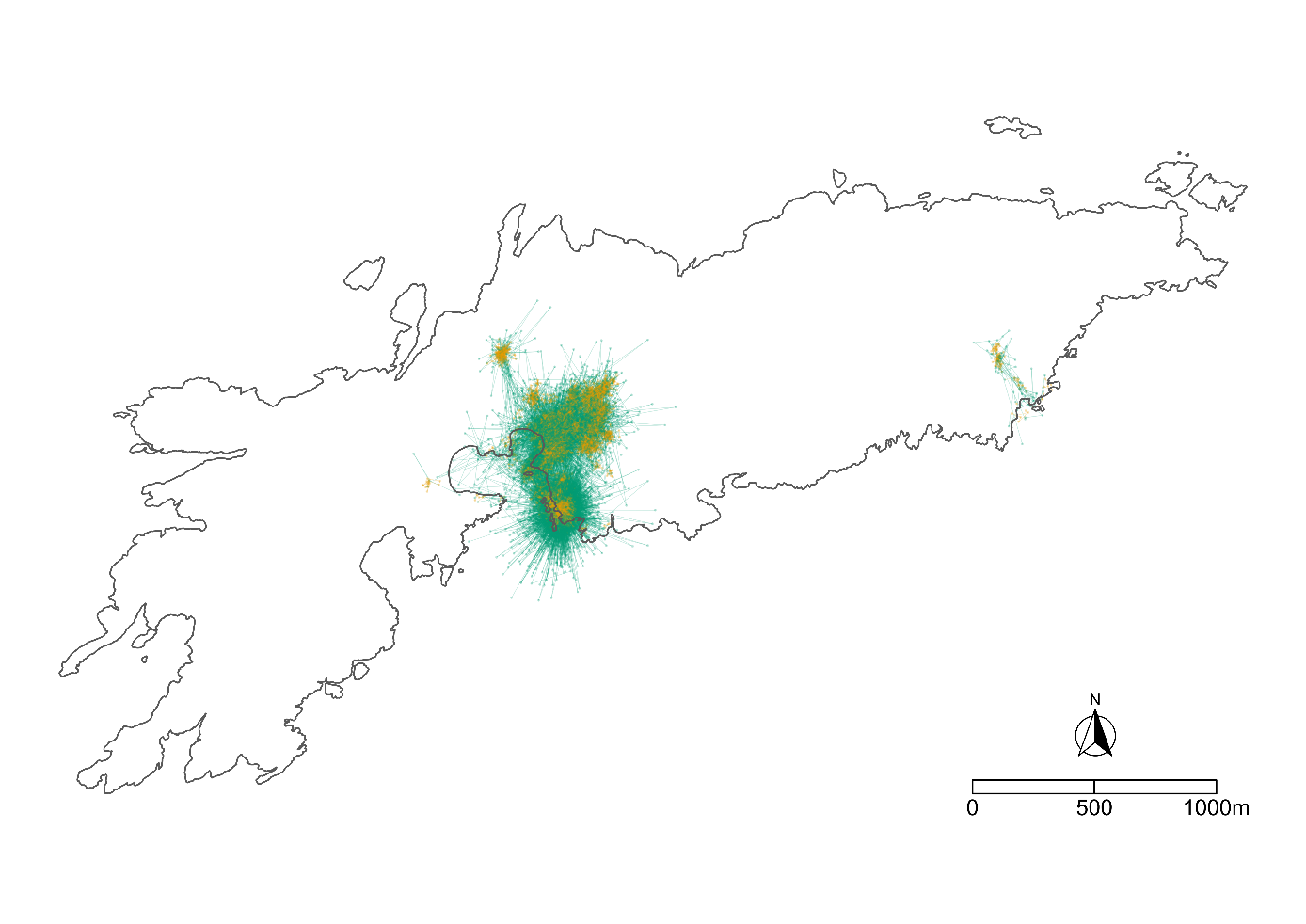

Figure S1: Map of Bird Island, South Georgia showing all recorded GPS positions and states recorded. The GPS tracks at the east end of the island are from a pup born at SSB and were confirmed with visual sightings of the individual at that location.

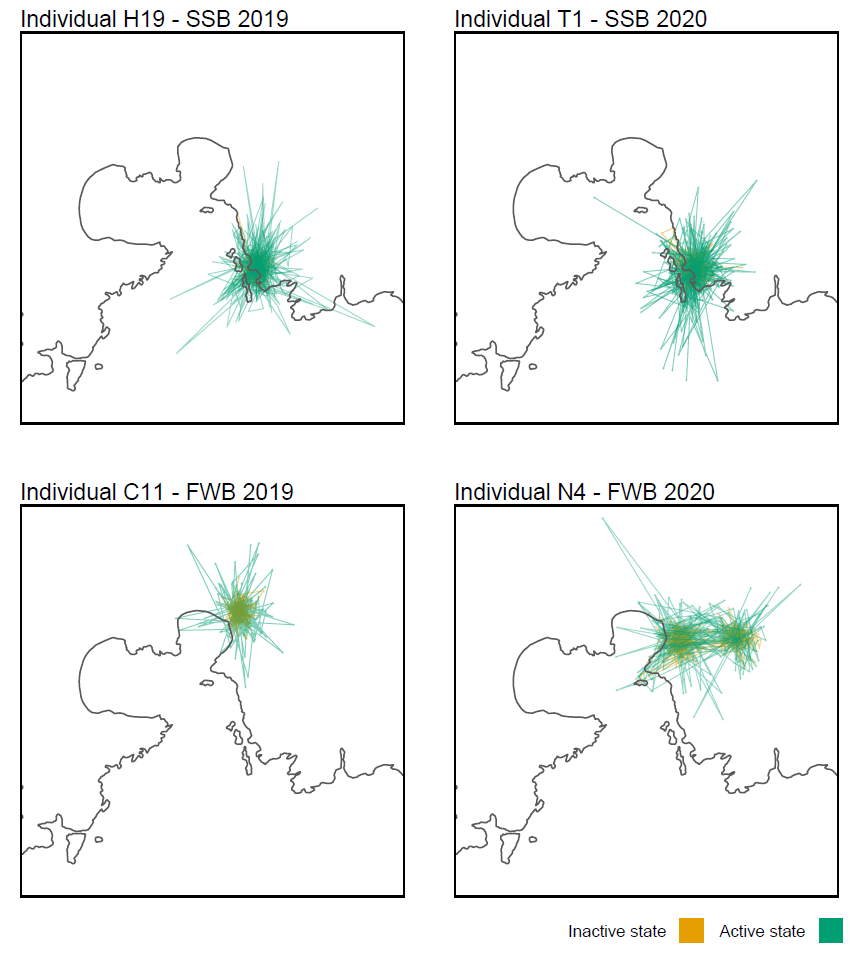

Figure S2: Representative examples of individual GPS tracks, including one pup from each breeding colony and breeding season.

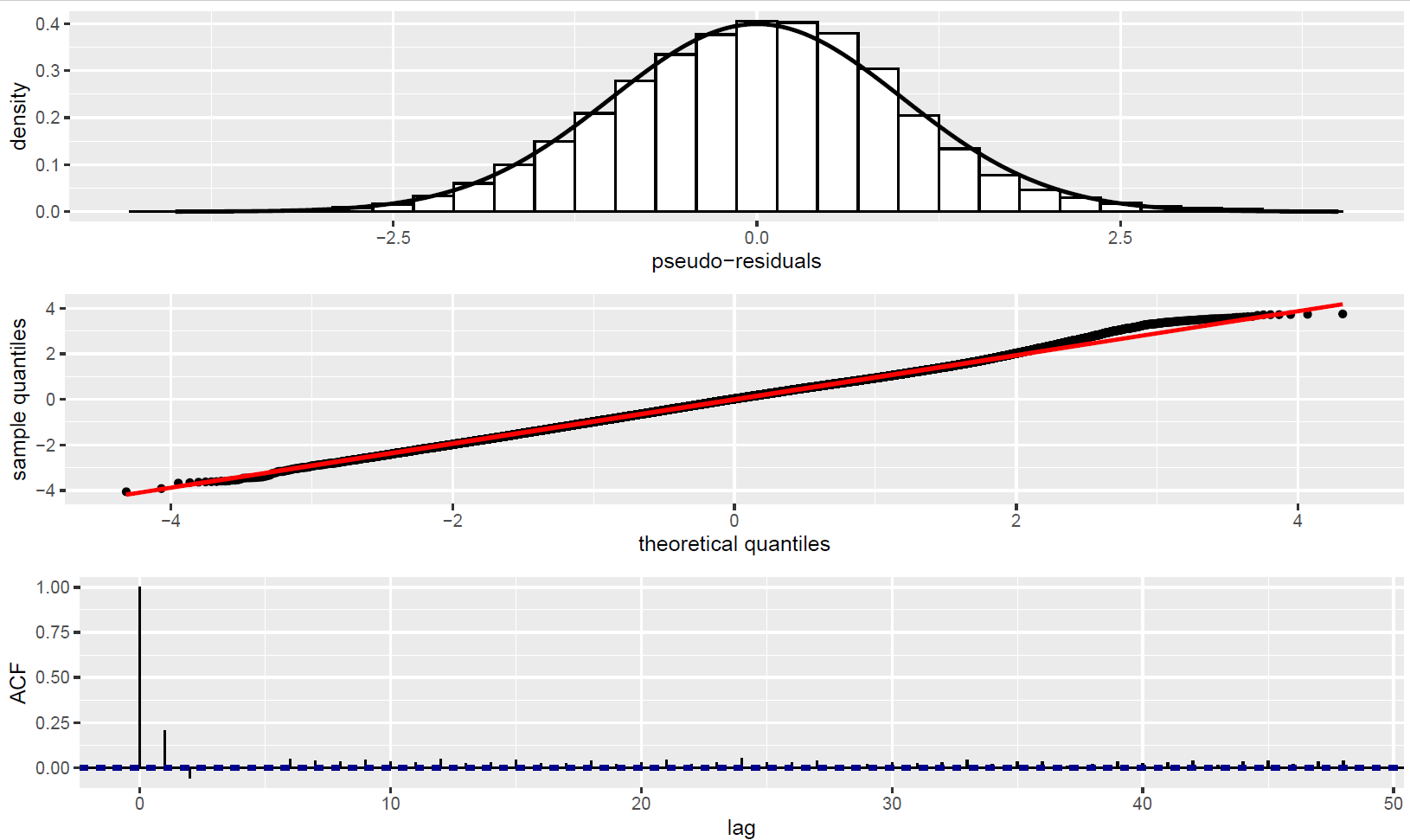
Figure S3: Pseudo-residuals of the final HMM.

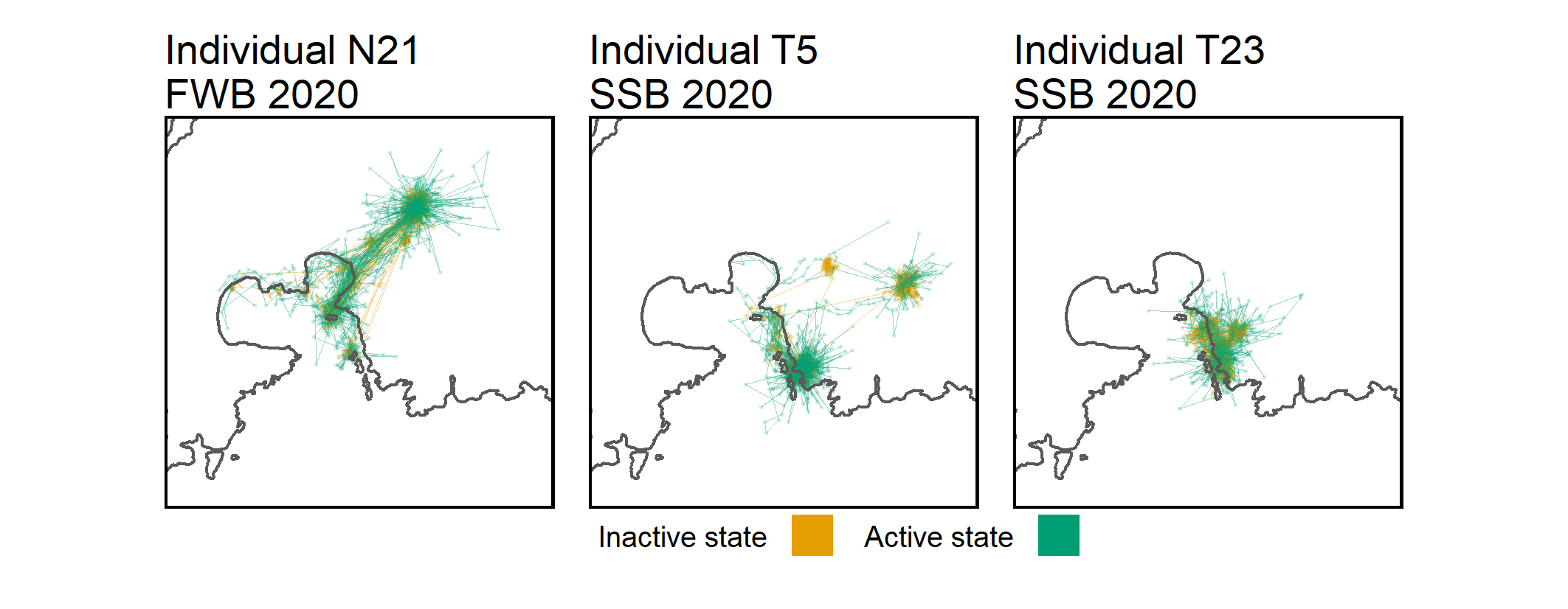

Figure S4: Five-minute interval GPS data were collected from three Antarctic fur seal pups, including at least one pup from each breeding colony. These data were gathered during the 2020 breeding season from pups between 30 – 50 days of age. Data were collected over a period of 13 – 19 days (mean = 3,965 data points). The figures show the decoded states from the HMM.

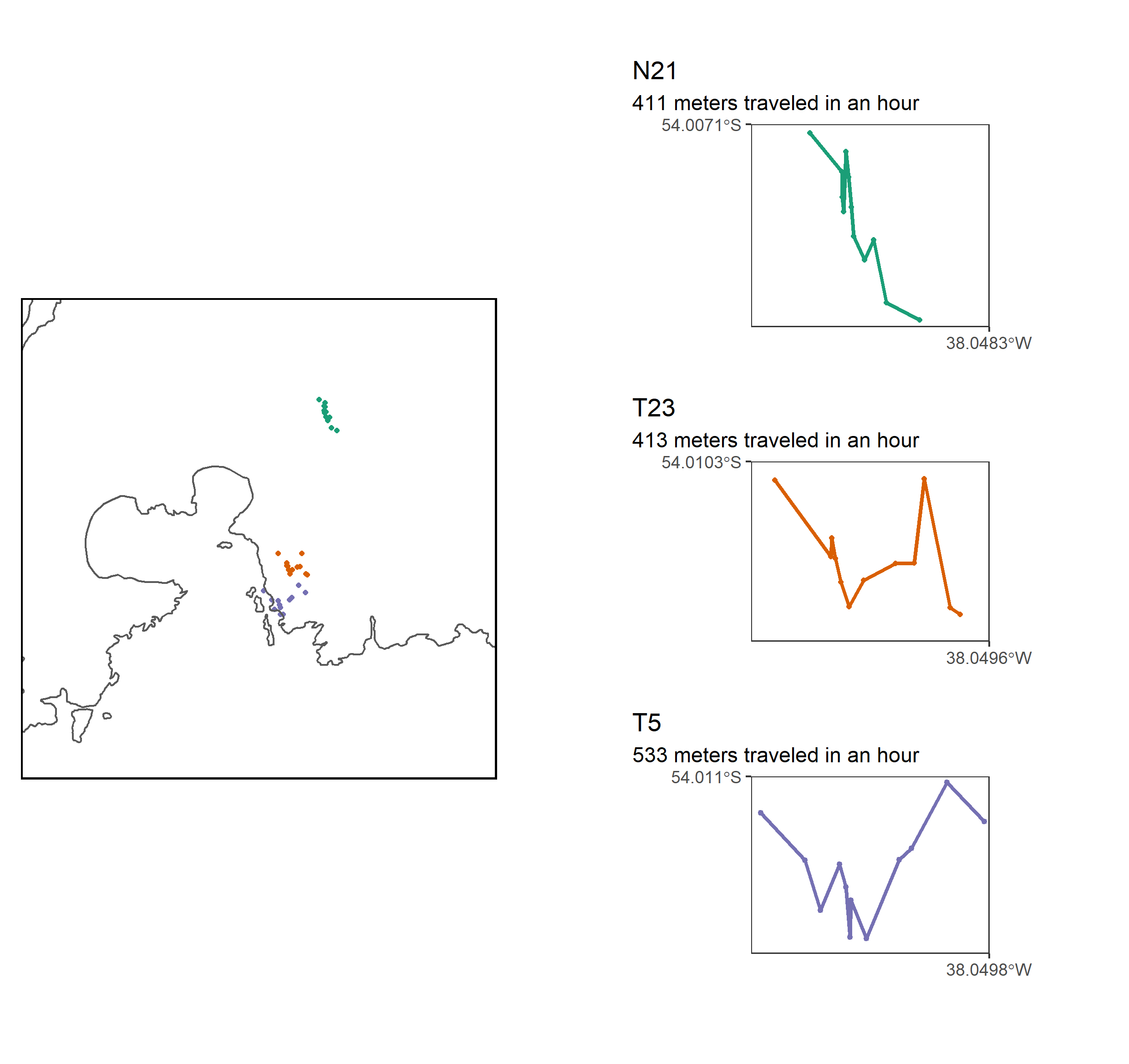

Figure S5: Five-minute interval GPS data from three pups (see Supplementary Fig. S4). Randomly selected examples are shown where a pup was inferred to have moved at least 400 meters in the space of one hour. All of these tracks reveal fairly consistent step sizes across the observation time window rather than a single, erroneously large step. Consequently, we believe that 400 meters is an appropriate step threshold for the HMM analysis.

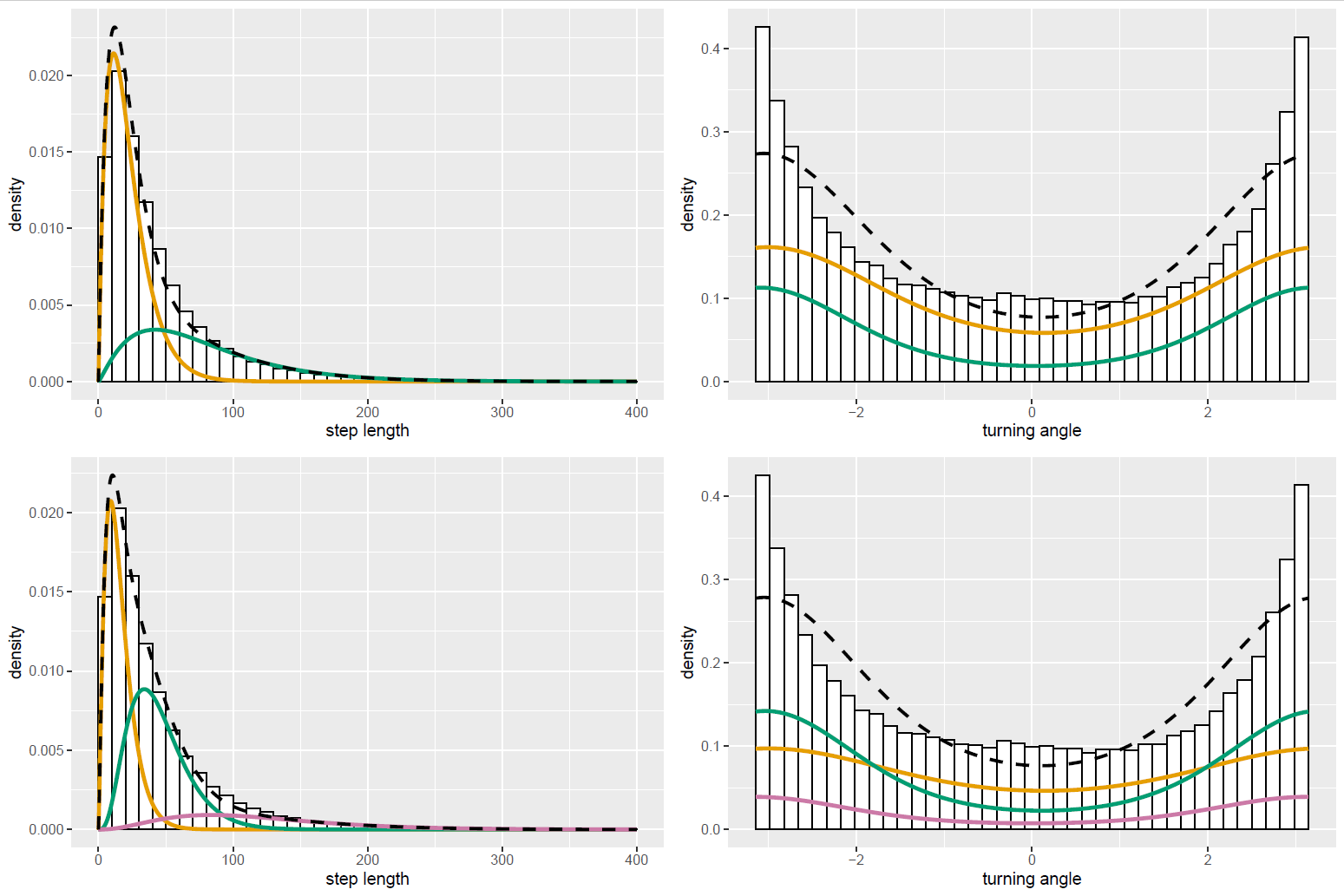

Figure S6: The number of states in an HMM must be specified before running the model and only models with a biologically meaningful numbers of states should be tested. We expected Antarctic fur seal pup movement to include a resting state (short step lengths), an active state (long step lengths with more directed movement), and potentially a local exploration state (moderate step lengths with high degrees of turning). Thus, we evaluated HMMs with two and three movement states. We also compared univariate HMMs to bivariate HMMs that included the turning angle between consecutive GPS locations. The marginal distribution under the fitted model and the empirical distribution is shown. In the 3-state HMM, the states’ distributions of both the step length and especially the turning angle overlapped greatly and did not allow us to adequately distinguish the active from the local exploration state. The distribution of the turning angle in the 2-state bivariate HMM was the same for both states and therefore did not add any information to the model. Overall, neither the additional state (three rather than two) nor the inclusion of the turning angle resulted in a significant improvement of model fit. Consequently, we restricted our final analysis to a parsimonious univariate HMM with two states.

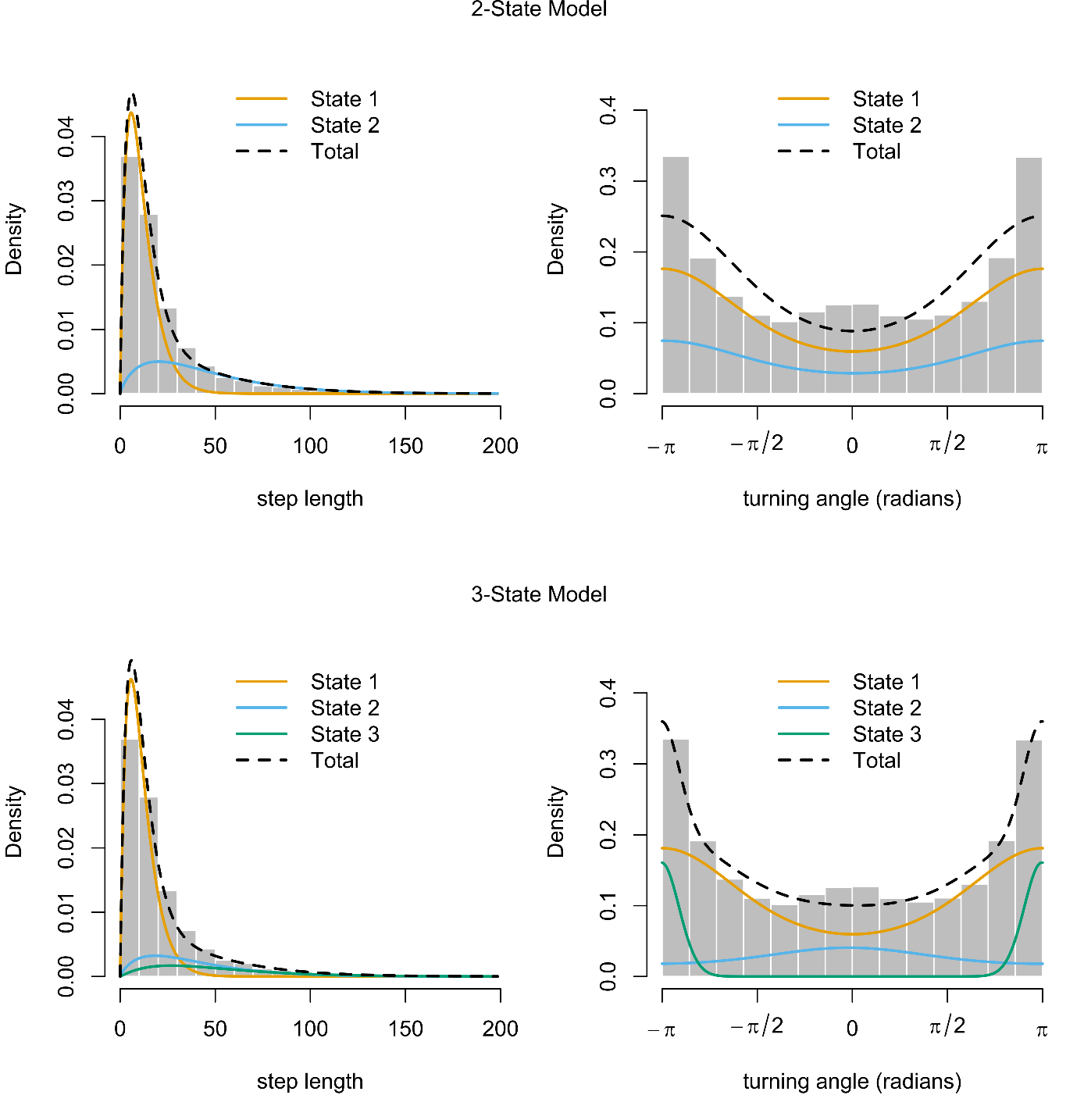

Figure S7: HMMs fitted to five-minute interval GPS data collected from three fur seal pups (see Supplementary Fig. S4). We evaluated HMMs with two and three movement states. With respect to the aims of our study, there was no relevant loss of information on pup activity using the coarser temporal resolution (hourly data).
